# Deep mutational scanning by FACS-sorting of encapsulated *E. coli* micro-colonies

**DOI:** 10.1101/274753

**Authors:** Lars Behrendt, Amelie Stein, Shiraz Ali Shah, Karsten Zengler, Søren J. Sørensen, Kresten Lindorff-Larsen, Jakob R. Winther

**Affiliations:** Linderstrøm-Lang Centre for Protein Science, Department of Biology, University of Copenhagen; Section for Microbiology, Department of Biology, University of Copenhagen; Division of Host-Microbe Systems & Therapeutics, Department of Pediatrics, University of California, San Diego; Center for Microbiome Innovation, University of California, San Diego; Present address: Environmental microfluidics group, Department of Civil, Environmental and Geomatic Engineering, Swiss Federal Institute of Technology

**Author notes:** Corresponding author: Linderstrøm-Lang Centre for Protein Science, Section for Biomolecular Sciences, Department of Biology, University of Copenhagen, Ole Maaloes Vej 5, DK-2200 Copenhagen N. These authors contributed equally (Lars Behrendt, Amelie Stein).

## Abstract

We present a method for high-throughput screening of protein variants where the signal is enhanced by micro-encapsulation of single cells into 20-30 μm agarose beads. Cells inside beads are propagated using standard agitation in liquid media and grow clonally into micro-colonies harboring several hundred bacteria. We have, as a proof-of-concept, analyzed random amino acid substitutions in the five C-terminal β-strands of the Green Fluorescent Protein (GFP). Starting from libraries of variants, each bead represents a clonal line of cells that can be separated by Fluorescence Activated Cell Sorting (FACS). Pools representing collections of individual variants with desired properties are subsequently analyzed by deep sequencing. Notably, the encapsulation approach described holds the potential for high-throughput analysis of systems where the fluorescence signal from a single cell is insufficient for detection. Fusion to GFP, or use of fluorogenic substrates, allows coupling protein levels or activity to sequence for a wide range of proteins. Here we analyzed more than 10,000 individual variants to gauge the effect of mutations on GFP-fluorescence. In the mutated region, we observed virtually all amino acid substitutions that are accessible by single nucleotide exchange. Lastly, we assessed the performance of biophysical protein stability predictors, FoldX and Rosetta, in predicting the outcome of the experiment. Both tools display good performance on average, suggesting that loss of thermodynamic stability is a key mechanism for the observed variation of the mutants. This, in turn, suggests that deep mutational scanning datasets may be used to more efficiently fine-tune such predictors, especially for mutations poorly covered by current biophysical data.

## Introduction

A detailed mechanistic and predictive understanding of the relationship between amino acid sequence and protein structure, stability and function impacts a vast number of practical and theoretical facets of biology and biotechnology. The design and engineering of proteins is commonly performed via targeted mutagenesis followed by a screening effort, resulting in the enrichment of desirable protein properties (1). Conventional screening approaches can address the (positive or negative) effects of a small number of mutations with high fidelity, yet are often drastically limited in the number of mutants that can be investigated effectively. Single-cell analysis by Fluorescence Activated Cell Sorting (FACS) is an attractive alternative that offers a high throughput as well as low (per sample) cost; however, it requires a fluorescence readout at the single-cell level. Likewise, microfluidic approaches tend to be labor intensive and require specialized and sometimes custom-designed equipment (2).

The combination of an efficient screening or selection system with next-generation-sequencing technologies allows the rapid accumulation of vast numbers of DNA sequences, and results in large amounts of data linking nucleic acid sequence to protein function (3). An interesting aspect of this is that both desirable and undesirable mutations can be sequenced, which allows for assessment of the accessible protein fitness landscape. Such *deep mutational scanning* (DMS) (4) has been implemented in a number of cases (5–8) and opened the possibility for detailed analysis of epistatic effects in protein sequences (9).

DMS is often combined with a system for genetic selection to allow for sorting variants with respect to the desired phenotype (e.g. catalytic function or binding). Such assays have, however, only been established for a limited set of systems (10). Likewise, biochemical assays that, due to lack of sensitivity require a microtiter format, are not accessible to DMS in the absence of costly robotic platforms.

Here we addressed the gap in sensitivity between single-cell screenings and standard microtiter formats by combining three well-established technologies into a novel high-throughput platform for the phenotypic screening of protein function (Fig. 1A). The approach comprises: (i) Microencapsulation, which allows for confinement of individual (fluorescent) bacterial cells into agarose microbeads. This encapsulation results in clonal amplification via cell growth within each microbead and hence a significant increase in fluorescence compared to single cells. (ii) Sorting of populations with specific phenotypes by FACS which is used as input material in (iii) parallel sequencing, which allows for the accumulation of DNA sequences at very high throughput. This procedure enables the genotypic probing of population subsets that exhibited specific fluorescence phenotypes. As a proof-of-concept, we applied this stepwise methodology on encapsulated micro-colonies of *E. coli* producing a library of GFP variants. Fusion to GFP would allow variant screening using our method for a wide range of proteins, and has successfully been used in similar applications (11).

**Figure 1.**
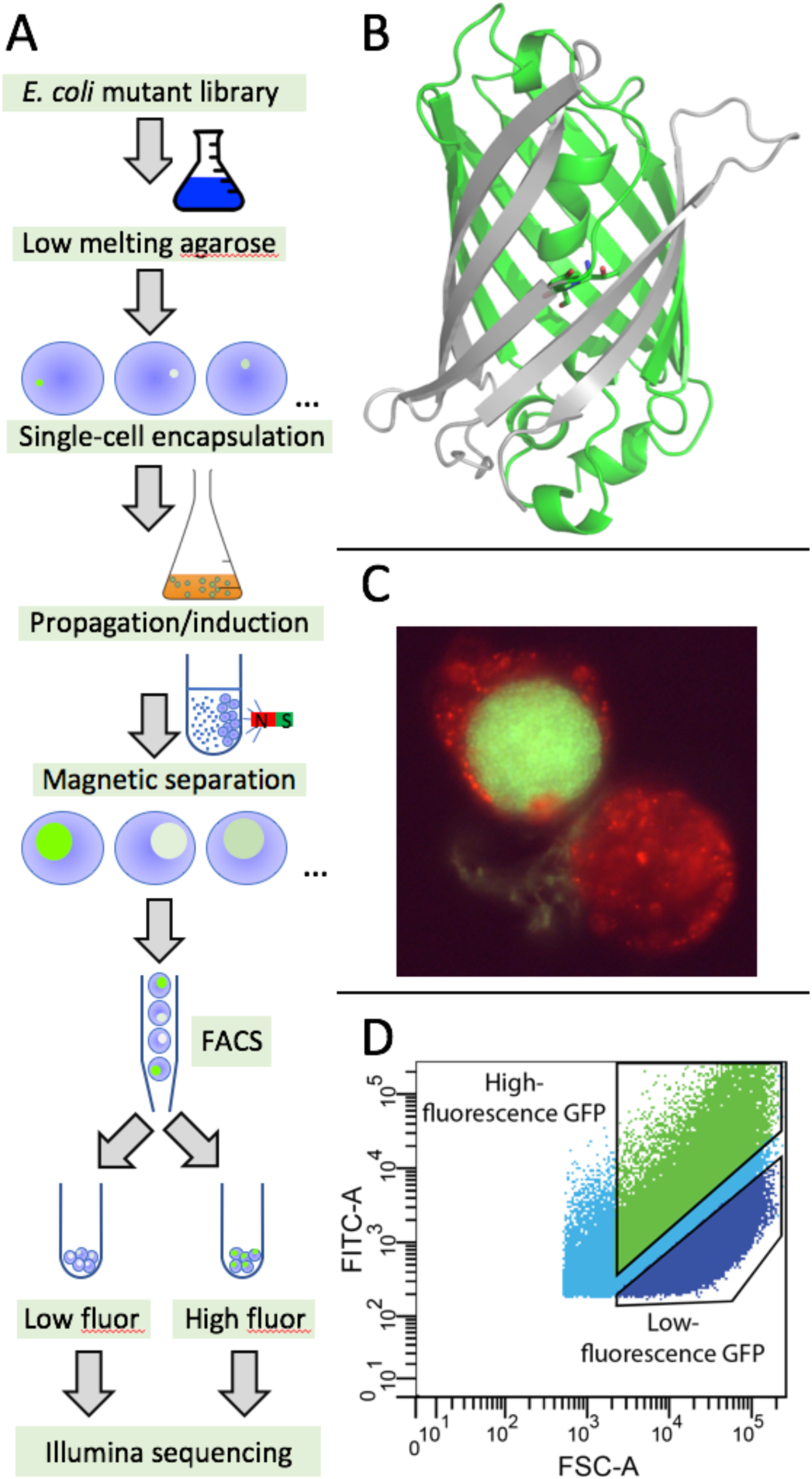
Micro-encapsulation and fluorescence activated cell sorting of GFP-expressing cells. (A) Experimental workflow. *E. coli* cells transformed with a mutant library are mixed at an appropriate dilution with low-melting agarose. Emulsification of this solution (including magnetic micro-particles) occurs by mixing agarose with mineral oil. Upon cooling, beads are solidified and contain <2 cells (shown as small green spheres) per bead. Cells are then grown inside beads using a conventional shaker flask with LB medium, to obtain micro-colonies.Beads are separated from free (planktonic) cells in a magnetic field and are subsequently sorted, using FACS, according to their fluorescence phenotype. The sorted populations are subjected to Illumina sequencing. (B) Structure of eGFP showing the five mutagenized β-strands in grey. The central chromophore is shown as sticks. (C) Agarose microbeads stained with a co-embedded red fluorescent dye for better visualization of the agarose gel. The upper left bead contains an *E. coli* micro-colony producing wild-type eGFP. The bead in the lower right does not contain bacterial cells. (D) FACS gating strategy employed to sort microbeads with fluorescent bacterial cells. Forward scatter (x-axis) was used as a proxy for colony sizes within microbeads and GFP intensity (y-axis) as a measure of fluorescence emission upon laser excitation. Dark blue and green represent the dim and bright fluorescent subpopulations, respectively.

## Results and Discussion

Our goal was to develop a bead-encapsulation method for signal amplification, and test its suitability for high-throughput screening of micro-colonies containing eGFP-protein variants. The key steps are outlined in Fig. 1.

### Mutant library

We chose GFP as a model system, as it allows for convenient distinction between folded (functional) and misfolded (dysfunctional) variants by means of measuring fluorescence. The exception are variants that affect chromophore formation. We thus aimed to assess the relationship between sequence and function on a large scale for a region of the GFP sequence that did not include the chromophore, and which spans adjacent β-strands in the three-dimensional structure. To obtain maximal sequence accuracy, we ensured that the mutated region was covered by Illumina sequencing (typically 300 base pairs in a single read), so that 5’ and 3’ reads would overlap significantly. However, it should be noted that this is not a requirement for the method. We chose to mutagenize the last third of the GFP open reading frame from amino acid residue 148 to residue 230 (Fig. 1B), which encodes five adjacent C-terminal beta-strands of the eGFP variant (12).

The mutagenized library was custom-synthesized by a commercial provider (BaseClear, Leiden, NL) and inserted into an expression vector under control of the *lac* promoter. Following library generation, optimal plasmid concentrations (∼one plasmid per successful transformant) were determined by titration (data not shown). The library of transformed *E. coli* cells was plated on selective media and grown over night, after which ∼50,000 transformed clones were washed off and appropriately diluted before proceeding to subsequent encapsulation.

### Encapsulation

We encapsulated cells in spherical agarose beads of FACS compatible sizes (20-30 µm diameter) using an emulsion-based approach (13). Encapsulated cells were grown over night at 37°C in LB growth medium containing antibiotics and expression was induced with IPTG; under conditions similar to the normal bacterial propagation. For visualization purposes, a red fluorescent color was added to the agarose during emulsification. Appropriate dilution of cells was visually confirmed using fluorescence microscopy and ensured the occurrence of microbeads with predominantly 0 or 1 cell within each bead. The observed fluorescent brightness in each microbead is expected to be proportional to the amount of (clonal) cells. As seen in Fig. 1C, a large number of cells can accumulate in a microbead. Although we have not experimentally assayed the number of cells per bead, we can make an estimate: If we assume that a collection of cells (as the one shown in Fig. 1) is spherical with about half the diameter of a 20 μm bead, the volume will be 4/3*π*r^3^ = 4/3*3.14*5^3^ μm^3^=4187 μm^3^. We assume a volume of 4 micron^3^ (14) for an *E. coli* cell grown on rich media. An irregular packing of spheres has a density of about 60% (15). To a first approximation, this is probably a reasonable estimate for *E. coli* cells as well. This gives us a number of cells per bead of 0.6*4200/4 ≈ 600 cells per bead.

During growth of cells inside microbeads, some cells may, however, escape encapsulation and are observed as single (planktonic) cells in the growth media. To avoid spill-over of cells into subsequent sorting steps we incorporated magnetic micro-particles into the agarose beads which allowed for repeated (magnet-assisted) washing steps, effectively removing most single cells from the bulk agarose beads.

### FACS-sorting of agarose encapsulated micro-colonies

Encapsulated clonal cell populations were sorted based on their relative fluorescence-emission intensity using FACS. Different sizes of cell clusters were microscopically observed in microbeads, and fluorescence intensity is thus expected to not only depend on the input plasmid but also on the size of the colony within the microbead. We addressed this issue by using forward-scatter (FSC) to normalize for size differences between cell-clusters, permitting us to assume that the fluorescence emission of similar sized cell-clusters is then predominantly protein-variant dependent. A total of 47,000 microbeads were sorted, with 18,600 originating from the gate with brightly fluorescing microbeads and 28,400 from the gate with dim fluorescing microbeads (Fig. 1D, Fig. 2).Correct sorting was visually verified on a small sample using fluorescence microscopy. Pools of sorted microbeads were used as input material for subsequent sequencing.

**Figure 2.**
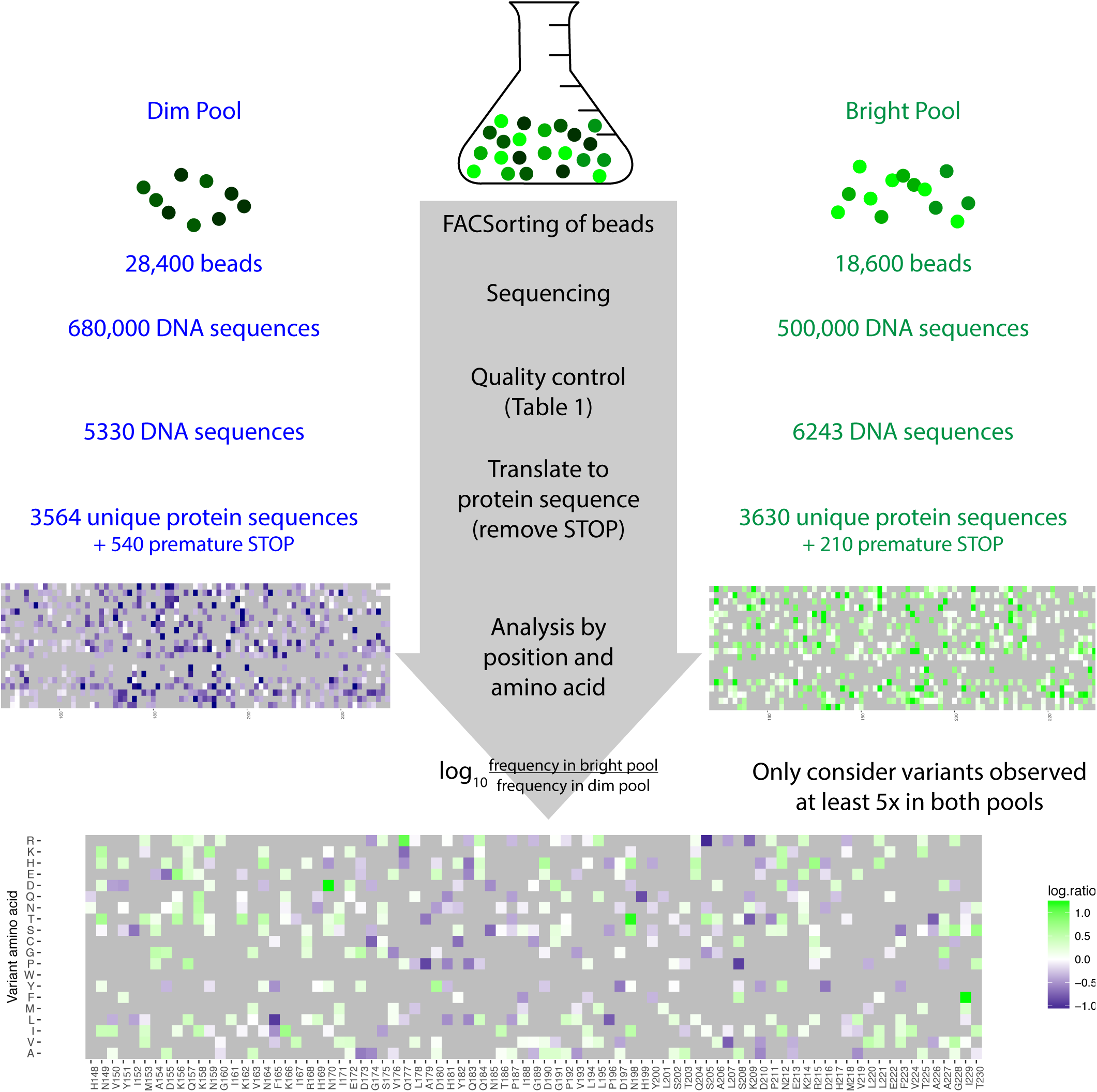
Overview of the sorting and quality control process from microbeads to interpretation of protein-level consequences. For details see main text. The purple-gradient (left) and green-gradient (right) matrices illustrate missense variants observed in the dim and bright pool, respectively, with darker colors indicating higher frequency, and gray indicating variants for which no data is available. The bottom center matrix illustrates the log_10_ ratio of the frequencies in the bright and dim pool, requiring at least 5 absolute occurrences in each pool. White indicates near-neutral variants (distribution in SI Fig. 4), green enrichment in the bright pool, and purple enrichment in the dim pool. Axes are the same for all three matrices.

### Sequencing

In order to link mutations to fluorescence phenotypes, we determined the underlying changes in nucleic acid sequences. This was done by preparing DNA from sorted beads as input material for PCR amplifications. Primers were designed to bind to within non-mutagenized regions of the sequence and, in combination, generated amplification products of correct sizes (405 bp). Primer designs contained additional overhangs to facilitate subsequent Illumina sequencing. A total of 1.2 x 10^6^ sequences were generated, of which the bright and dim pools of fluorescent libraries contributed 5.0 x 10^5^ and 6.8 × 10^5^ sequences, respectively.

### Assembly of DNA sequences and analysis of mutations

Compared to conventional next-generation-sequencing, where mostly unique DNA sequences are mapped against a corresponding reference genome, the extraction of a large number of nearly identical sequences poses different challenges, as mutations must be distinguished from sequencing error with high fidelity (16). The number of beads sorted by the FACS provided us with a maximum possible number of 18,600 brightly and 28,400 dimly fluorescent variants (Fig. 1D, Fig. 2). The number of unique raw DNA sequences, however, outnumbered these limits by more than an order of magnitude, indicating a need for rigorous quality control to remove erroneous sequences.

To this end, paired-end sequence files corresponding to FACS-sorted populations were mapped against the DNA sequence of the original eGFP gene in a multiple sequence alignment. Next, for the portion of the eGFP gene sequence where the read pairs overlapped, differences between pairs were assumed to result from sequencing errors, while agreement with each other was assumed to comprise a mutation. In addition, we required that the variant DNA sequence appeared in at least 2 independent pairs of overlapping reads. The number of unique sequences recovered in this manner was ∼30,000 from each sorted population (Table 1), and still considerably greater than the number of originally sorted beads in the bright pool. Thus, even with identical sequences in the overlap region and at least two reads showing the same sequence, sequencing error was a major problem, which, given typical NGS reliability, is not entirely unexpected (16). This was also indicated from calculation of the apparent number of mutations for the different number of reads. Accordingly, reads occurring at a low frequency (2-3 times) appeared to be substantially more prone to mutation than those with more than 4 reads (Table 1). As mutation rates should be independent of the number of reads, we chose to use this metric for estimating the number of identical reads needed to validate a sequence as genuine. The overall frequency of mutations for 5 or more reads was 1.63% for the dim fraction and 1.24% for the bright fraction. The observation that mutations are less frequent in the bright pool is expected.

**Table 1:** Read-depth and mutation frequency within bright and dim subpopulations. The number of reads indicate how many times a given DNA sequence occurred. “No. of sequences” indicates number of unique sequences with the specified number of reads. The mutation frequency indicates the number of base substitutions relative to wild type. It is likely that the higher “mutation rate” seen with very few reads (2 or 3) reflects coincidental sequencing errors.

|  | Dim |  | Bright |  |
| --- | --- | --- | --- | --- |
| No. of reads | No. of sequences | Mutation frequency (%) | No. of sequences | Mutation frequency (%) |
| 2 | 28949 | 2.16 | 32025 | 1.66 |
| 3 | 4805 | 1.99 | 6044 | 1.49 |
| 4 | 1680 | 1.80 | 2058 | 1.40 |
| 5 | 917 | 1.71 | 1039 | 1.34 |
| 6 | 603 | 1.63 | 623 | 1.30 |
| 7 | 421 | 1.60 | 434 | 1.30 |
| 8 | 360 | 1.60 | 306 | 1.30 |
| 9 | 302 | 1.46 | 240 | 1.27 |
| 10 | 223 | 1.63 | 245 | 1.21 |
| 11 | 194 | 1.45 | 182 | 1.22 |
| 12 | 137 | 1.64 | 168 | 1.23 |
| >12 | 2173 | 1.64 | 3186 | 1.19 |
| **At least 5** | **5330** | **1.63** | **6423** | **1.24** |
| Total | 40764 |  | 46550 |  |

Overall the library of sequences resulting from the >4 reads cut-off represents a fairly broad distribution of base substitutions (SI Table 1); transitions and transversions were 0.49% and 0.75%, respectively, for the bright population and 0.56% and 1.06%, respectively for the dim population.

### Effects of mutations at the protein level

Full-length DNA sequences containing mutations with a read-coverage >4 were translated into protein sequences. Unique DNA sequences in dim and bright populations (Table 1) collapse into 3,564 and 3,630 unique protein sequences with an average of 2.9 and 2 amino acid substitutions across the 83 residues of the mutagenized region, respectively. Variants leading to premature STOP codons were removed. Given the diversity of the resulting sequences, and in absence of systematic data on double-, triple-mutants etc., we performed a position-based analysis based on the cumulative mutation counts observed in the dim and bright pools. The subsequent analysis focuses on point mutations at the protein level. Specifically, we calculated the relative frequency of each individual amino acid substitution in the bright vs. dim population (Fig. 2). Given the experimental setup, this means that each individual variant was typically observed in the context of 1-2 additional (random) mutations on the protein level. In order to increase robustness and minimize the influence of errors and noise (e.g. encapsulation of two different clones in the same bead, sorting uncertainties, or integration over different sequence contexts), we only performed this calculation for amino acid substitutions that we observed in at least five distinct protein contexts (i.e., unique protein sequences) in each pool. Like with the acceptance of error rate in DNA sequencing, the number of minimum independent occurrences represents a compromise. With this threshold, 503 of the 507 possible single-nucleotide changes leading to amino acid substitutions were observed in the bright and dim populations. Ninety single-nucleotide-variants were only observed in the dim population, indicating that these are most likely unstable. Four variants were only observed in the bright populations and thus likely stable (I161V, K162R, K166R, P211A). Four variants were not observed in either population (V150G, I161N, H199R, K209E), which may be due to screening depth, library coverage, or post-sequencing quality control as described above. In addition to one double-nucleotide change, S202Y (Fig. 2), the remaining 409 single-nucleotide variants were observed in both populations.

### Analysis of effects of amino acid substitutions

We next set out to analyze the effects that variants have on GFP structure and function, and to assess whether biophysical calculations could predict them.

First, we analyzed silent mutations, which are unlikely to affect function once the protein is made, and may thus serve as a baseline for background mutation frequency between the two pools. As expected, the bright pool on average contained more synonymous changes than the dim pool, though for individual positions the dim pool contained more silent mutations (SI Fig. 2). We took the 90^th^ percentile of the log_10_ ratio of silent mutations (0.29) as a significance threshold for the interpretation of non-synonymous changes (SI Fig. 3).

**Figure 3.**
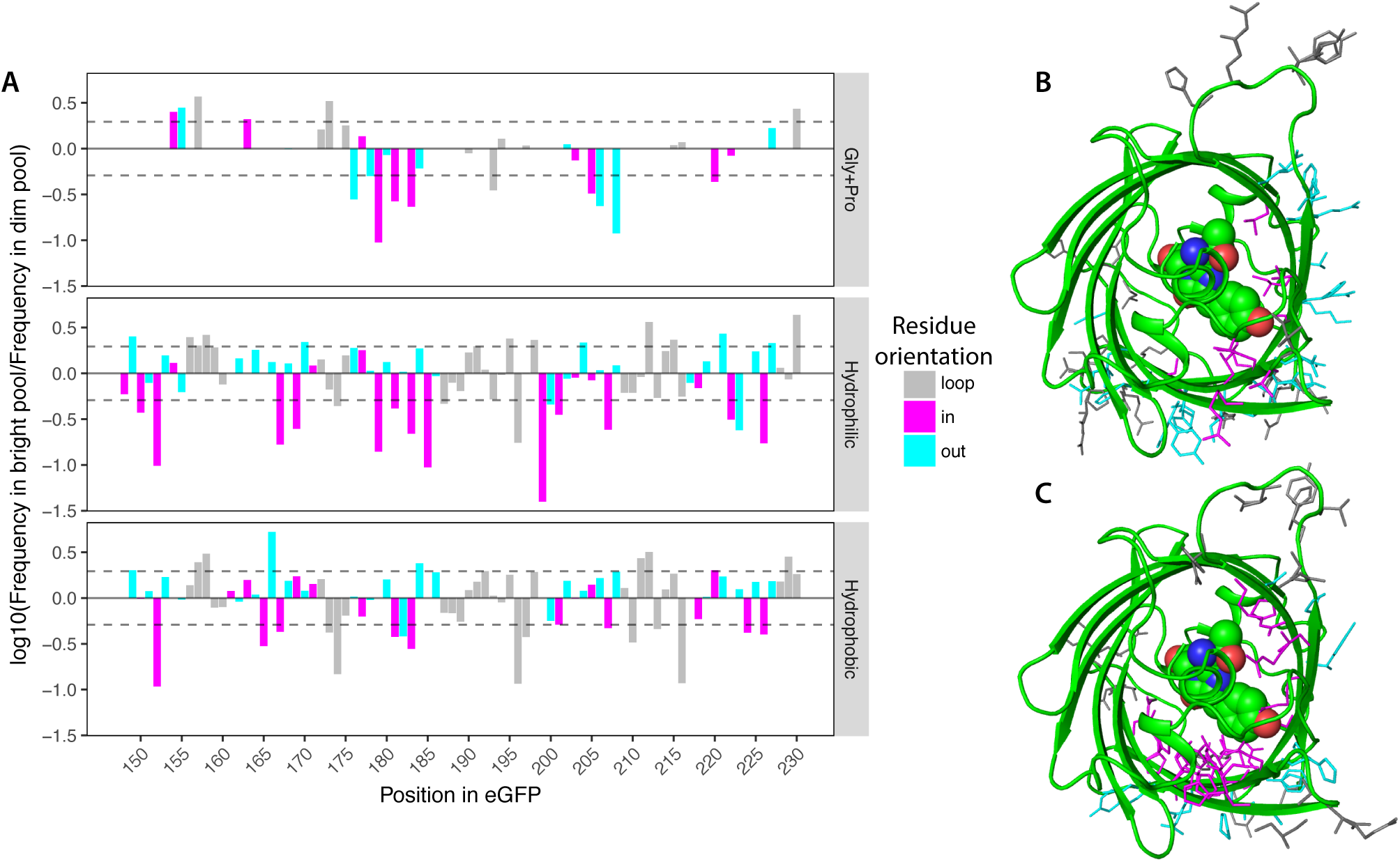
Mutations introducing hydrophilic residues facing inward are strongly depleted in the pool. (A) Analysis of enrichment and depletion by the orientation of the residue in native GFP (PDB: 1EMA, (19); magenta, pointing in; cyan, pointing out; gray, loop) and hydrophobicity of the mutant residue (hydrophobic residues, A/C/F/I/L/M/V/Y/W; hydrophilic, D/E/H/K/N/Q/R/S/T). Dashed lines indicate the significance threshold, based on synonymous changes observed in the bright and dim pools. (B) Enriched and (C) depleted variants, respectively, mapped onto the structure of eGFP using the same color scheme as in panel A. Note the large number of magenta (hydrophilic) residues pointing inwards in the depleted variants in panel C.

For a missense mutation, the *a priori* expectation would be that amino acid residues pointing into the core of the β-barrel should be more restricted than those pointing out, and that positions with native, buried hydrophobic residues would be particularly constrained (17). Indeed, the strongest signal we observed in our assay was a selection against mutations pointing into the GFP molecule, especially for mutations to hydrophilic residues (Fig. 3). In fact, 3 of the 4 single-nucleotide changes not observed in either population point into the GFP core and may therefore be detrimental. Among the observed variants, at 14 positions pointing into the β-barrel, mutations were significantly depleted. In contrast, this was only the case for 6 positions that point into solvent, and 9 positions in loops. Further, only 18 positions revealed significantly enriched variants, and 9 of these were found in loops (Fig. 3). This agrees with recent analyses showing more deleterious variants in sheets and helices than in loops (17). Nevertheless, the overall mutational robustness in the loops here is still fairly low, in line with another recent analysis which found highest robustness in helices (18). Generally, as illustrated in Figures 3 and 4, enrichment of non-synonymous mutations beyond the significance threshold is rare.

**Figure 4.**
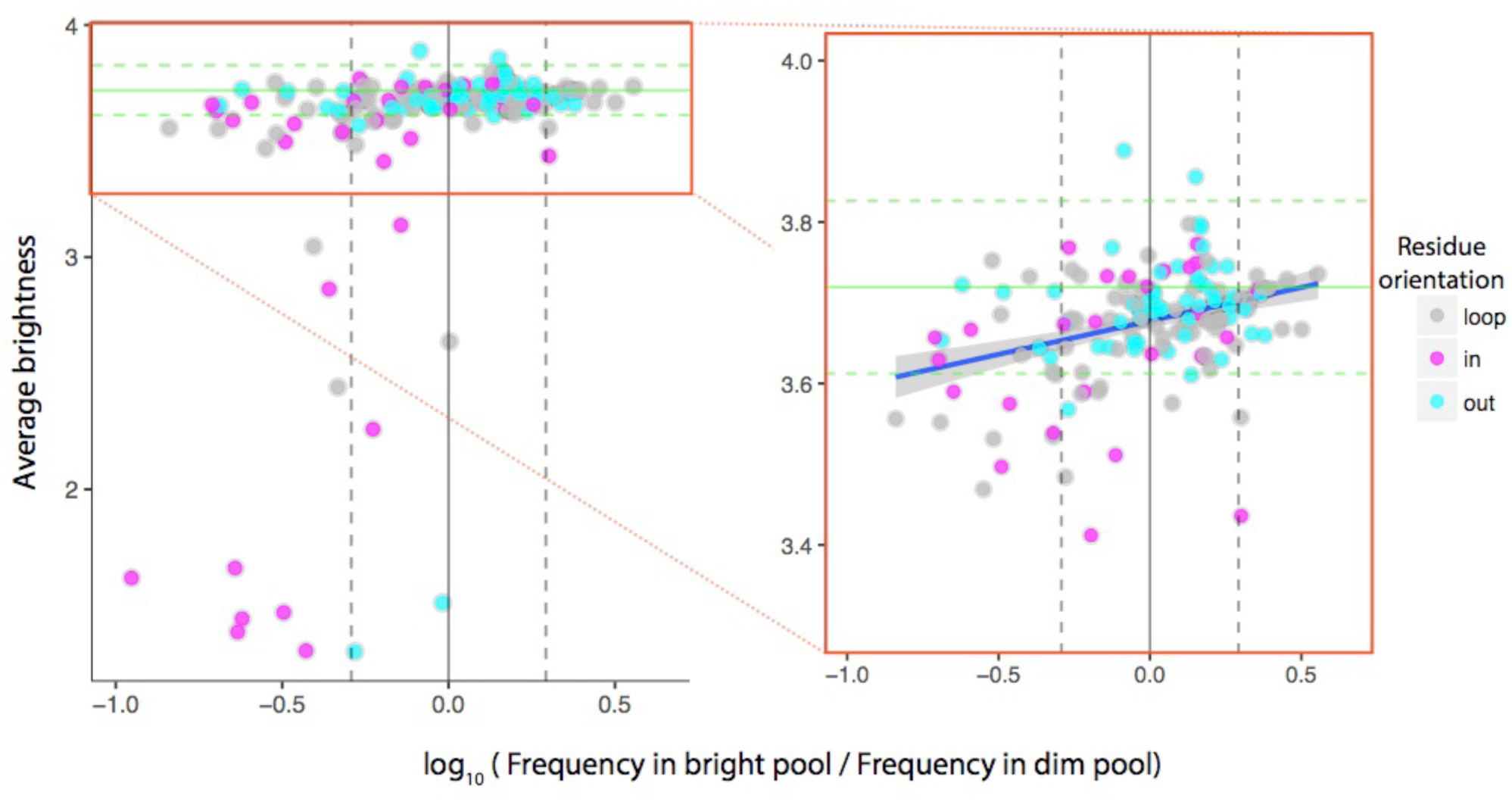
Comparison of the enrichments observed in our study versus a previous one based on FACS sorting of single cells. (7). X-axis as in Fig. 3, y-axis shows average brightness for each position and residue group calculated from the published data. Dashed vertical lines indicate the significance threshold, based on synonymous substitutions. Horizontal lines indicate wild-type GFP brightness ± one standard deviation. The best agreement between the two high-throughput approaches is demonstrated for mutations at positions pointing inwards (magenta, correlation coefficient 0.56). Most of the severely detrimental variants (lower left corner) also point into the molecule. Overall though, most tested variants are scored as near-neutral by both methods.

We then compared the enrichment observed in our data to another recent large-scale GFP mutational study (7). There is surprisingly little overlap between the two datasets. This may, however, be explained, as the two studies used different starting DNA sequences, and the vast majority of variants observed in both studies were those reachable with single-nucleotide changes from the respective starting sequence. We thus focused on the 287 mutations that were quantified in both deep mutational scans. For comparison purposes, we grouped variants by orientation and amino acid category like in Fig. 3 and calculated the average brightness within each group (Fig. 4).

The Spearman correlation coefficient for all pairs is 0.43, however when only considering residues that point into the molecule, it is 0.56 (Fig. 4). As discussed above, residues pointing inwards are expected to be the most constrained, which is captured by both studies. Overall, both experimental approaches classify most of the assessed mutations to be close to neutral, which is in line with other studies on mutational tolerance in proteins (17). The distribution of fitness scores for the two methods is rather different, with a narrow peak close to wild-type fluorescence for the previously published data vs. the broader distribution of log-ratios we calculated (Supplementary Fig. S4). This difference is also reflected in the almost binary fluorescence distribution observed in the previously published study of GFP (7) vs. the continuous and overlapping distributions of our data (Supplementary Fig. S5). This may arise from the signal amplification provided by the microbeads used in this study. Generally, the agreement between the two studies is comparable to that between other high-throughput screens (20).

### Predictive capability of biophysical stability calculations

Lastly, we performed biophysical stability calculations for the mutations assessed in this study, and compared them to our screening-based log ratios described above. Predictions of change in stability (ΔΔG) are widely used in biotechnology (22) including protein design (6), but also in the assessment of non-functional or disease-associated protein variants (23, 24). We used the established tools Rosetta (25) and FoldX (26) to calculate ΔΔGs for each of the possible single amino acid changes in our mutagenized section of GFP (SI Figs 6 and 7).

The average Rosetta ΔΔGs for each enrichment category (neutral, enriched, depleted) agreed well with our experimental classification (Fig. 5), and the predicted ΔΔG classified most enriched and depleted variants correctly as shown by an area under the curve (AUC) of 0.83 in a receiver operating characteristic (ROC) analysis (Fig. 5A). However, we also observed a number of outliers, including predictions of enriched mutations as being destabilized and predictions of depleted mutations as having near-wild-type stability (Fig. 5B).Averages for enriched vs. neutral variants were only substantially different for mutations pointing into the molecule, while many detrimental mutations were correctly recognized regardless of orientation (Fig. 5C). The FoldX ΔΔG calculations painted a very similar picture (SI Fig. 8). This is in agreement with previous observations that destabilizing mutations are predicted more accurately than stabilizing changes (27). Overall the agreement between our experimental readout and the ΔΔG calculations indicates that changes in stability are likely major molecular reason underlying the change in phenotype (GFP fluorescence) we observe. Some deviation from biophysical calculations might be explained by variation at the nucleotide level which can also lead to changes in expression levels, which would similarly affect the experimental readout. However, previous analyses of ΔΔG prediction performance have similarly revealed many outliers despite good agreement on average (28), indicating considerable room for improvement. We suggest that one underlying issue may be that current biophysical predictors are trained on a dataset consisting mainly of hydrophobic truncations; primarily to alanine (Fig. 6A) (25, 26, 29, 30). In contrast, the mutations assessed here were dominated by amino acid substitutions that are in adjacent codons (Fig. 6B). These include many small-to-large amino acid substitutions, which were extremely scarce during training of predictors, and which may require more sophisticated protocols that allow for backbone adjustment (25, 31).

**Figure 5.**
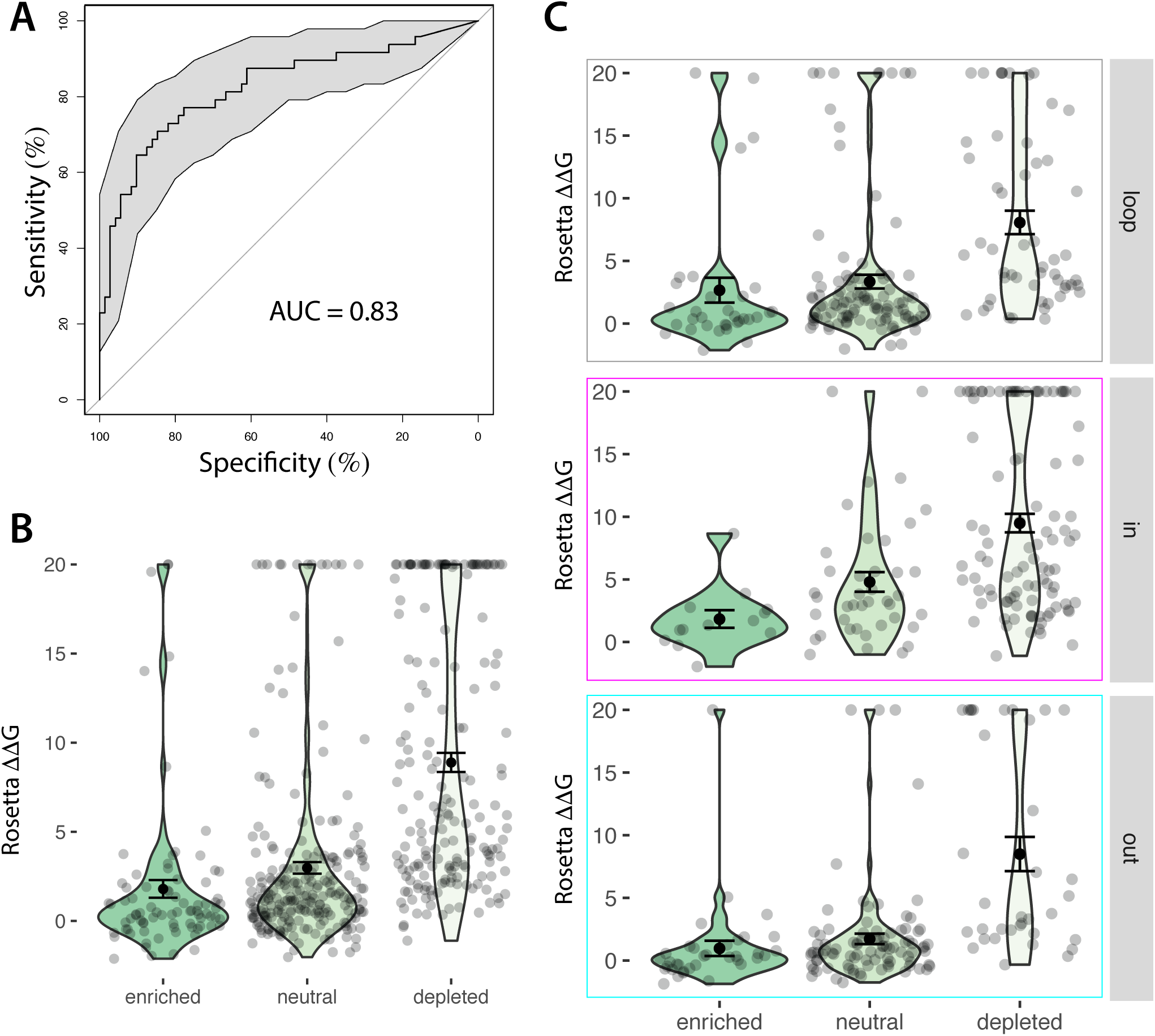
Comparison of experimental readout with computational prediction of stability. (A) Receiver operating characteristic (ROC) analysis of the performance of Rosetta ΔΔGs in classifying GFP variants as tolerated (enriched) or not (depleted) (21). (B) Rosetta ΔΔG predictions for all mutations assessed in our screen, separated by enrichment category. “Neutral” is defined based on synonymous changes observed in our dataset. Rosetta predictions were carried out as described previously (see Methods). (C) Analysis of ΔΔG predictions split by residue orientation shows that discrimination of enriched vs. neutral changes is only significant for residues pointing into the β-barrel.

**Figure 6.**
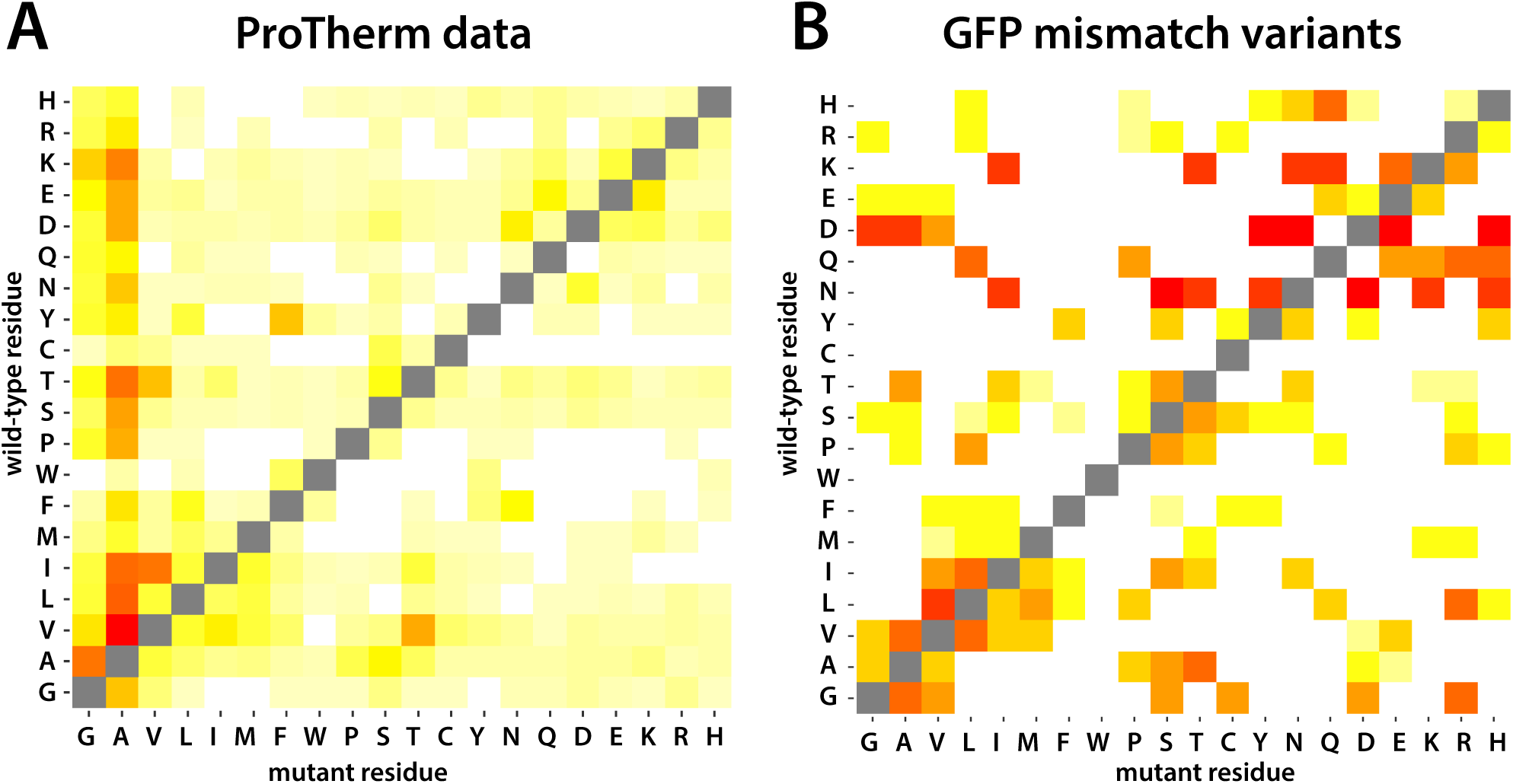
Maps of amino acid substitutions used for training computational methods vs those observed in the present study. (A) Single residue mutations for which biophysical measurements are available in ProTherm, aggregated from hundreds of individual studies (29). (B) Single residue mutations observed in this study. Red tiles indicate the most common changes, orange and yellow are less common, white tiles indicate missing residue pairs.

## Concluding remarks

We have established a method that uses microbead encapsulation to perform DMS for protein variant phenotypes. As mentioned above, it is relevant to compare this approach to microtiter formats and microfluidic techniques used in cases where an enzymatic assay is required. Due to the clonal signal-amplification, this method is well suited for screening efforts that target low-fluorescing variants or require cell propagation, and may also provide an increased dynamic range. Beyond the generation of sequence-function maps for the specific protein mutagenized in the assay, cumulative data from DMS studies may provide valuable data and insights for protein sequence-stability or –function maps in general (17). Such data may in turn be used to benchmark computational methods for assessing the impact of mutations on proteins (32), or indeed to train new prediction methods (33). With the development of generic assays that effectively capture also the effects of protein stability (11, 34), we envision that deep mutational scanning data may serve to improve existing but – as illustrated above – far from perfect biophysical calculation programs. Indeed, a first DMS-based predictor of missense mutation consequences (33) shows good overall performance, and integration with mechanistic biophysical approaches is likely to increase both predictive power and accuracy. The increased dynamic range of the presented microbead approach could thus allow for the separation of subtle differences in stability and for the prospective fine-tuning of such predictors. Additional future applications include the use of encapsulated cells that could be exposed to (or co-embedded with) fluorogenic substrate prior to FACS; aiding in the elucidation of phenotype-functional relationships of natural enzymes and for high-throughput screening in enzyme design.

## Materials and Methods

### Construction of pFH2102::S65TGFP mutagenesis libraries

Random mutagenesis was performed on a 250 bp subset on the 3’ end of the S65T-GFP gene (NCBI accession number 20473140). Mutations were introduced by a commercial provider (BaseClear, Leiden, The Netherlands) in a *de-novo* synthesis approach. Using fixed primers at non-mutagenized sites, the mutagenized GFP gene was amplified and then cloned into pFH2102 using XhoI and EcoRI restriction sites.

### Plasmid titration and cloning

The pFH2102::S65TGFP mutagenesis library was diluted to concentrations ranging from 5 x 10^−8^ g – 5 x 10^−13^ g in 10x dilution steps. Electrocompetent GeneHog cells (Invitrogen, Carlsbad, CA, USA) were transformed with 1 µl of the diluted plasmid pool and electroporated using a Bio-Rad Micropulser according to the manufacturer’s instructions. After electroporation cells were immediately resuspended in pre-warmed SOC media and left to grow at 37°C for 1h at 300 rpm. Transformed cells were then plated onto pre-warmed LB plates with 100 µg ml^−1^ Ampicillin and left to grow overnight at 37°C. Three technical replicates were plated for each of these reactions and colony forming units (CFU) at given plasmid concentrations were enumerated after 18h of growth. Subsequent transformations were calibrated to ensure that the majority of transformants had only one plasmid per cell. The plasmid titration resulted in an optimal plasmid concentration of 2.5 pg per transformation and was determined based on the midpoint of the linearly increasing part of the CFU vs. plasmid concentration curve.

Using this optimal pFH2102::S65TGFP mutagenesis library concentration GeneHog cells were transformed using the above protocol. Approximately 50.000 individual colonies were washed off the selective plates using a Drigalski-spatula and pre-warmed LB medium containing 100 µg ml^−1^ Ampicillin. These cells were used, after appropriate dilution, for encapsulation in agarose.

### Agarose-encapsulation

For encapsulation into agarose beads, 200 µl of the appropriately diluted cells were mixed with 1 ml of pre-heated (45°C) 1.5% SeaPlaque LMP agarose (Lonza, Rockland ME, USA), 20 µl magnetic beadMag particles (Chemicell, Germany) and then transferred into 15 ml warm (45°C) paraffin oil. Several of these reactions were done in parallel in order to obtain enough material for FACS sorting. The oil:agarose suspension was emulsified using a custom blender according to the following protocol: 2 min at 3000 rpm at room temperature, 1 min at 3000 rpm on ice and 6 minutes at 1400 rpm on ice. The resulting agarose beads were separated from the oil phase by centrifugation (2000 x g) and washed 3X with sterile PBS (pH 8.0). For a more comprehensive description of the encapsulation protocol please refer to (13).

The resulting beads were placed into a 250ml Erlenmeyer flask containing 100 ml of sterile LB augmented with 100 µg ml^−1^ Ampicillin and 1 mM IPTG and left to grow overnight under 37°C at 300 rpm. The next day the agarose beads were centrifuged at 2000x g for 5 minutes, washed once with fresh LB, pelleted and the supernatant removed. The pellet was resuspended in fresh LB and aliquoted into Eppendorf tubes placed on a magnetic Eppendorf rack (DynaMag, Thermo Fisher Scientific). The magnetic agarose beads were separated from the supernatant and repeatedly washed using sterile PBS and vortexing, thereby removing both planktonic bacteria and bacteria attached to the outside of beads. After completion of the washing procedure the beads were pooled and filtered through a custom made 30 µm nylon mesh (Frisenette ApS, Denmark) inserted into a Pop-top syringe filter (Whatman, UK).

### Mutant screening using Fluorescence-activated cell sorting

All flow cytometric analysis was performed on a FACSAria III (BD Biosciences, USA) using a 488-nm laser in conjunction with a GFP specific emission filter set (denoted FITC, 530/30nm bandpass) and forward scatter detection (FSC). The gating scheme to sort highly fluorescing beads from less fluorescing beads utilizes a size threshold (FSC=500) as well as GFP minimum fluorescence (FITC=200). This assures that only beads with cells expressing fluorescing GFP variants are sorted but implies that beads with non-fluorescing (e.g. stop-codons) cells are not. Exclusion gating was chosen empirically to avoid any potentially remaining planktonic cells (FITC:<200) and to include only agarose beads (FSC>500; Fig. 1D). The high-fluorescence (FITC: 500-10^5^ relative fluorescence units (RFUs)) and low-fluorescence gating (FITC: 200-10^4^ RFUs) was set so individual slopes were identical and thereby parallel for the size-dependent FSC.

A total of ∼47000 beads were sorted, of which ∼18500 contained highly fluorescing GFP variants and ∼28000 contained cells with lower fluorescing GFP variants. During preliminary experiments, we observed that the size of cell clusters within beads affects GFP emission intensity of the sorted beads. To compensate for this effect we employed FSC as a means to normalize for different cell cluster sizes and only performed sorting of beads within the same FSC gate. This approach relies on the assumption that cell growth and GFP expression underlies the same distribution in both brightly and dimly fluorescing populations.

### PCR amplification of inserts and sequencing

All sorted beads were centrifuged at 12.000 x g for 2 minutes. The supernatant was carefully removed until a residual volume of ∼40 µl was left and 12.5 µl of Lyse and Go PCR Reagent (Pierce Biotechnology, Rockford, IL, USA) was added. The beads were heated for 2 minutes at 95 °C and immediately placed on ice. In addition to cells sorted via FACS the original transformation library (GFPorig) was amplified to determine its inherent sequence diversity. All 50 µl PCR reactions were composed of either 10 µl of the lysed beads or 5 ng of library (GFPorig) as DNA template. The PCR reaction components were 1 µl of the primers GFP-dom-5911-Fv’ and GFP-dom-300-Rv’ (10 µM each, SI Fig 1), 0.02 U µl^−1^ Phusion HF-II DNA polymerase (Thermo Fisher Scientific, Hvidovre, Denmark), 1 µl dNTP’s (final concentration 200 µM each) and 10 µl 5X Phusion HF Buffer. PCR amplification used the following cycle conditions: 98 °C for 30s, 30 X (98 °C for 10s, 60 °C for 30s and 72 °C for 15s), final extension was done at 72 °C for 10 minutes and samples were hereafter kept at 4°C. The resulting PCR amplicons (= 405 bp) were visualized on a 1% agarose gel, the correct bands excised and purified using the Qiaex gel extraction kit (Qiagen Nordic, Copenhagen, Denmark). Based on the extracted DNA, a sequencing library was prepared according to the manufacturer’s instructions using the Nextera kit (Illumina Inc., San Diego, CA, USA). Library concentrations were quantified and checked using a fragment analyzer and paired-end (2 × 250 bp) sequenced on an Illumina MiSeq system (Illumina Inc., San Diego, CA, USA).

### Calculation of enrichment ratios

To assess whether variants on the protein level are favored or disfavored, we calculate the log_10_ of the relative frequency of each individual amino acid substitution in the bright pool vs. the dim pool. Numbers of synonymous variants(SI Fig. 2) and local log_10_ ratios (SI Fig. 3) fluctuate across the length of the mutagenized strands, likely due to base pair composition. We chose the 90^th^ percentile of the distribution, 0.29, as our threshold for significance. Variants with absolute enrichment or depletion below this threshold are considered near-neutral.

### Biophysical stability predictions

Rosetta ΔΔGs were calculated using a previously published protocol (25) (protocol 13), version with GitHub SHA-1 6922a68c56c0a3c5f64570c55097ba5d5439e22c (Nov 2016). A special parameter file for chromophore was generated based on PDB 1ema and the instructions in https://www.rosettacommons.org/docs/latest/rosetta_basics/preparation/preparing-ligands”. Command lines for the initial minimization and subsequent ΔΔGs calculations were: /path/to/rosetta/source/bin/minimize_with_cst.macosclangrelease -in:file:l lst -in:file:fullatom -extra_res_fa CRO.params - ignore_unrecognized_res -fa_max_dis 9.0 -database /path/to/rosetta/database/ -ddg::harmonic_ca_tether 0.5 - ddg::constraint_weight 1.0 -ddg::out_pdb_prefix min_cst -ddg::sc_min_only false > mincst.log; sh/path/to/rosetta/source/src/apps/public/ddg/convert_to_cst_file.sh mincst.log > input.cst

## Commands used to calculate ΔΔGs

/path/to/rosetta/source/bin/ddg_monomer.linuxgccrelease -database /path/to/rosetta /database -ex1 -ex2 -ddg:local_opt_only true -constraints::cst_file ../input.cst -ddg::min_cst true -s ../min_cst.GFP_het_Rosetta_renum_0001.pdb -ddg:iterations 32 - ddg:mut_only -resfile ../GFP_mutagenize.resfile -extra_res_fa ../CRO.params

To account for the large sampling space, 32 trials were performed for each variant. ΔΔGs reported here are calculated based on the mean energies of the 3 lowest-scoring replicas of each variant.

FoldX (26) ΔΔGs were calculated by first running RepairPDB on the starting structure, and then using the BuildModel command to generate each individual single residue variant. The chromophore is removed during RepairPDB. The protein backbone is kept constant by FoldX. After 5 iterations for each variant, the energy difference of the mean scores was reported.

